# Characteristics of putative keystones in the healthy adult human gut microbiome as determined by correlation network analysis

**DOI:** 10.1101/2023.11.20.567895

**Authors:** Franziska Bauchinger, David Seki, David Berry

## Abstract

Keystone species are thought to play a critical role in determining the structure and function of microbial communities. As they are important candidates for microbiome-targeted interventions, the identification and characterization of keystones is a pressing research goal. Both empirical as well as computational approaches to identify keystones have been proposed, and in particular correlation network analysis is frequently utilized to interrogate sequencing-based microbiome data. Here, we apply an established method for identifying putative keystone taxa in correlation networks. We develop a robust workflow for network construction and systematically evaluate the effects of taxonomic resolution on network properties and the identification of keystone taxa. We are able to identify correlation network keystone species and genera, but could not detect taxa with high keystone potential at lower taxonomic resolution. Based on the correlation patterns observed, we hypothesize that the identified putative keystone taxa have a stabilizing effect that is exerted on correlated taxa. Correlation network analysis further revealed subcommunities present in the dataset that are remarkably similar to previously described patterns. The interrogation of available metatranscriptomes also revealed distinct transcriptional states present in all putative keystone taxa.

**IMPORTANCE:** The work presented here contributes to the understanding of correlation network keystone taxa and sheds light on their potential ecological significance. By employing a robust workflow based on bootstrapping and subsampling techniques, we identify putative keystone species at the genus and species level. This emphasizes the importance of considering taxonomic resolution when investigating correlations. The potential impact of keystones on community stability provides valuable insights for systematic microbiome manipulation. Furthermore, the observed clusters of co-occurring taxa align well with recent findings and emphasize the reproducibility and relevance of the identified patterns in microbial community composition. We are able to add a functional dimension to the analysis with the identification of distinct transcriptional states in putative keystone taxa, highlighting their functional versatility and adaptability.

## INTRODUCTION

Microbiomes are characterized by diverse interspecific and microbe-environment interactions. The net outcome of microbial activities and interactions produces the observed community structure and function, and interactions are thought to be crucial for maintaining community stability and conferring resistance and resilience in the face of disturbance. Borrowing a concept from macro-ecology (1), the idea of keystone species has intrigued many microbiologists as a potential ecological mechanism that facilitates community stability and resilience. Microbial keystones, whether defined at a species level or a higher taxonomic grouping, are characterized as highly interconnected taxa that exert a strong influence on the entire community, shaping the microbiome irrespective of their abundance (2). Loss of a keystone taxon from the community would be expected to disrupt the microbiome and cause major shifts in community composition and function. Their crucial role in maintaining community structure is what makes keystone taxa a particularly interesting target for human gut microbiome research.

The gut microbiome is essential for human physiological function and health by performing many important services including metabolizing complex food molecules (3), training the immune system (4), and providing colonization resistance against pathogens (5). Identifying keystone taxa in the healthy gut microbiome and understanding their role in the community could provide us with novel microbiome-targeted intervention strategies for dysbiosis-related diseases such as inflammatory bowel disease, cardiovascular disease, and obesity (6). Some species have been proposed as keystones in the human gut microbiome based on empirical evidence, such as *Akkermansia muciniphila* (7) and *Christensenella minuta* (8). *A. muciniphila* was shown to facilitate the growth of butyrate producers by degrading host-derived mucosal sugars in a mucus-dependent cross-feeding community (7). *C. minuta* showed protective properties against diet-induced obesity in mouse models and modulated the intestinal microbiota in an *in vitro* model inoculated with fecal samples from obese individuals (8). However, the experimental identification of keystone taxa in the human gut microbiome is challenging as intervention studies are difficult to conduct and animal and *in vitro* models can have limited translatability for the human gut microbiome. Consequently, researchers have moved towards bioinformatics and data analysis to identify putative keystone taxa (9–11).

A popular approach to identify keystone taxa in microbiomes is to use sequencing-based microbial abundance profiles to construct and analyze correlation or co-occurrence networks (12). The assumption is that positive or negative interspecific interactions, be they direct or indirect, lead to positive or negative correlations between the abundances of the respective taxa. It has been shown by Berry and Widder (13) that in simulated communities certain network features, namely node degree, closeness centrality, betweenness centrality and transitivity, are indicative of a taxon’s keystone potential and can be used to identify keystone taxa with 85% accuracy.

Several studies have identified putative keystone taxa based on correlation network analysis (10, 14, 15) and proposed them as targets for subsequent experimental studies. However, we still lack an understanding of what ecological features these keystones share or which functional niches they occupy that lead to their observed characteristics in correlation networks. We can speculate that a keystone taxon provides a conserved, specific function in the gut microbiome that is not provided by other taxa and is therefore crucial in maintaining community structure and function. Alternatively, keystone taxa might be functionally versatile and able to fill available ecological niches, thereby providing the needed functional redundancy and resilience to microbial communities. One can also imagine a mixture of both functions, either provided by a single taxon or a keystone guild.

With this study, we aim to further our understanding of putative keystone taxa in the human gut microbiome. We establish a workflow to construct robust and statistically significant correlation networks with FastSpar (16) and identify correlation network keystones based on their network features, as proposed by Berry and Widder (13). We then utilize this workflow to interrogate publicly available metagenomes and metatranscriptomes from large human gut microbiome studies. We find that detection of network keystones is highly sensitive to taxonomic resolution and are able to identify putative keystone taxa only at species and genus level. Correlation network analysis further suggests a community stabilizing effect of putative keystone taxa and reveals co-occurring subcommunities present across gut microbiomes.

## RESULTS

### Establishing a workflow to build significant and robust correlation networks

In this study, we established a workflow that results in correlation networks based on both highly significant as well as robust correlations (Fig 1). A major concern when constructing correlation networks is the inherent compositionality of community abundance profiles derived from sequencing data, which can lead to spurious correlations. We utilized FastSpar (16), a C++ implementation of the well established tool SparCC (17), which identifies significant correlations and excludes spurious correlations based on a bootstrapping algorithm. We constructed correlation networks with FastSpar on subsamples of our dataset and subsequently constructed a consensus network based only on correlations that were identified as significant in at least 20% of the individually constructed networks. This workflow ensures that correlations kept in the consensus network are representative of both the entire dataset as well as subsamples. We used this procedure to identify taxa that robustly exhibit correlation network keystone features in the gut microbiomes studied.

**FIG 1.**
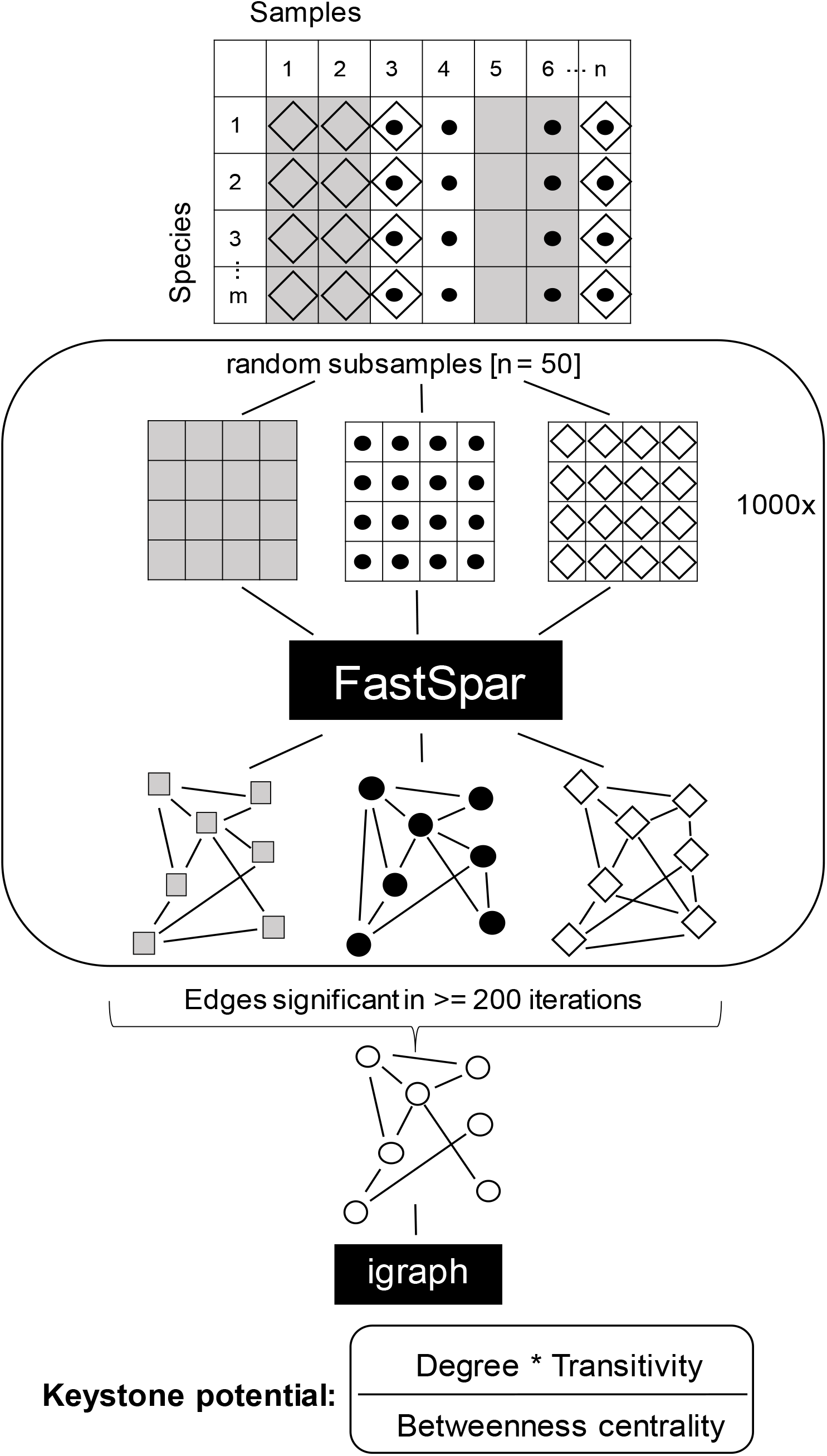
Workflow for the construction of robust and significant correlation networks. The dataset is subsampled 1,000 times and correlation networks are constructed from subsamples using FastSpar. Correlations identified as significant in >= 200 iterations are used to construct a consensus network. The mean correlation strength of all significant correlations is used as the correlation strength for the consensus network. Igraph is used to calculate relative node degree, transitivity and betweenness centrality.

### Community structure and species properties

In order to evaluate keystone taxa in the healthy human gut microbiome, we leveraged two large studies of healthy human adults, which included 580 paired metagenome and metatranscriptome samples from 131 individuals (18, 19). We first estimated whether the community compositions of the two datasets were similar enough to be combined by performing an ordination analysis and computing the similarity between communities on a species level. This revealed that while samples from the two datasets are somewhat separated, they still exhibit a strong overlap (Fig S1A). To ensure statistical independence between samples we randomly selected one sample per participant, resulting in an even larger overlap between the two datasets (Fig S1B). To further confirm the validity of combining datasets, we investigated the site similarity of the included samples by computing the overall Jaccard similarity between the communities (presence and absence of species), as suggested by Berry and Widder (13). Overall, the studied communities exhibit a similarity of 36%, exceeding the suggested lower limit of 20%. These results confirmed the suitability of combining the datasets for computing correlation networks and estimating keystone potential. To gain a better understanding of the data used for this study, we analyzed relative abundance, prevalence and transcriptional contribution of prevalent species (Fig S1C). We observed a large variance in relative abundance, both within and between species. As expected, several species of *Bacteroides*, such as *Bacteroides vulgatus, Bacteroides uniformis* and *Bacteroides stercoris*, as well as *Faecalibacterium prausnitzii* and *Prevotella copri* are highly abundant in the dataset. More abundant species also tend to contribute more strongly to metatranscriptomes. This could reflect actual contribution to transcription, but may also be a result of better annotation quality in reference genomes from highly abundant species. While many highly abundant species are also highly prevalent across the dataset, there are multiple exceptions. Most notable and well known is the low prevalence of *P. copri,* a species that has previously been observed to be abundant in certain gut microbiomes while mostly absent in many others, particularly in cohorts from Europe and North America (20). Focusing on taxa exhibiting a prevalence of at least 20% across all studied samples, we then constructed correlation networks as described above.

### Correlation networks reveal a loss of structure and keystone potential at lower taxonomic resolution

Previous work has shown that closely related taxa can form networks to utilize complex substrates in the human gut (21), but stable interactions between distantly related taxa have been also observed (22). It is, however, poorly understood how taxonomic resolution affects correlation analysis and specifically the identification of putative keystones. We therefore computed correlation networks at different taxonomic resolutions (order, family, genus and species) and investigated their structure. Overall, networks showed reduced modularity with increased taxonomic resolution (genus and species), while cohesion decreased from species- to order-level networks (Fig 2A). Modularity measures the extent to which a given network is divided into modules, with a higher modularity indicating greater divisions within the network. In contrast, the cohesion of a network estimates the minimum number of nodes that need to be removed to result in a weakly connected network, and a higher cohesion therefore indicates a more tightly connected network. Constructing correlation networks at low taxonomic resolution produced weakly connected networks while networks constructed at genus- and species-level are more cohesive and less modular. This observation is further confirmed by the observed distributions of network features, namely relative node degree (ND), transitivity (T) and betweenness centrality (BC). The relative degree of a node (= a taxon) is the number of edges a node has (correlations to another node) relative to the size of the network (total number of nodes). Transitivity indicates whether all nodes correlated with a given node are in turn correlated. The betweenness centrality of a node estimates how many shortest paths between two nodes (smallest number of edges connecting two nodes within a network) go through this given node. Correlation network keystones are characterized by high node degree, high transitivity and low betweenness centrality (13). The probability density functions of these network features, particularly transitivity and betweenness centrality, are flatter at lower taxonomic resolution (Fig 2B). In the node degree distribution we observe a slight flattening as well as a general shift towards a higher degree at lower taxonomic resolution. Flatter probability density functions, and therefore a more equal distribution of network features across all taxa, suggest that any one taxon is less likely to exhibit a high keystone potential. We confirm this by computing the keystone potential, using the formula *keystone potential = (ND*T)/BC*, and comparing it across taxonomic resolutions (Fig 2C). As expected, we observe very low keystone potential at family and order level, indicating that such broad taxonomic groups are unlikely to exhibit structuring effects on the overall community. In contrast, we see both genera and species with a high potential to act as correlation network keystones. Interestingly, the pattern of keystone potential is not consistent from genus to species, with some taxa exhibiting a higher potential at genus level and seemingly losing it at species level and vice versa. In summary, these results suggest that correlation network keystones are solely found within genera or species, and we thus focused our downstream analyses on these two taxonomic levels.

**FIG 2.**
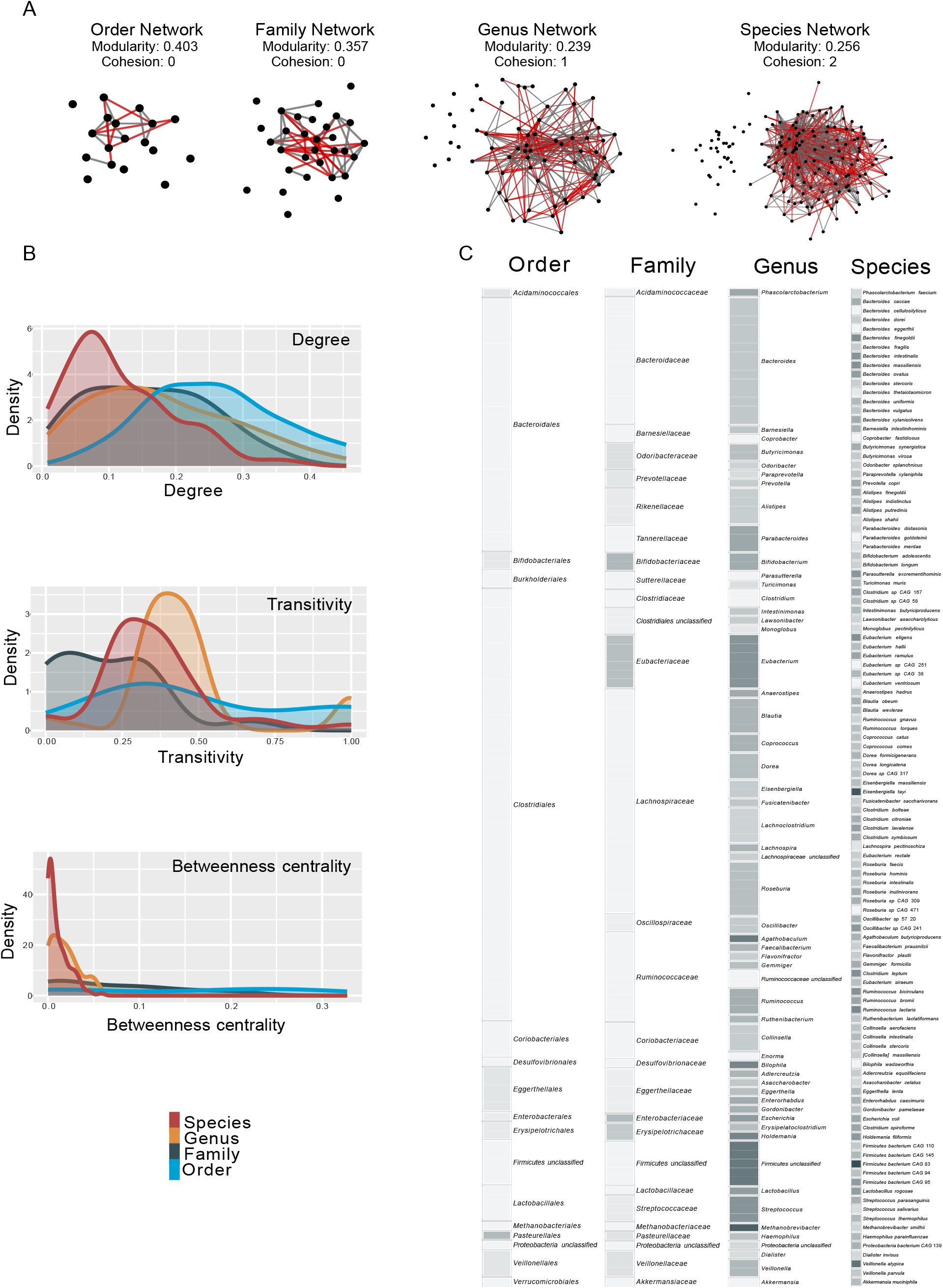
Distribution of correlation network features and keystone potential. (**A**) Correlation networks constructed on order, family, genus and species level. Network nodes (black circles) indicate taxa, network edges connecting the circles indicate positive (grey) and negative (red) correlations between taxa. (**B**) Density functions of relative node degree (relative to total network size), transitivity and betweenness centrality of taxa within a correlation network, coloured by taxonomic resolution. (**C**) Keystone potential of taxa present in >= 20% of analyzed samples. The keystone potential of each taxon is indicated as a coloured square, with white indicating very low keystone potential and dark gray indicating high keystone potential.

### Few taxa have a relatively high keystone potential

We next analyzed the distribution of keystone potential in genera and species to better understand the network properties of correlation network keystones. Both on genus as well as on species level the distribution of keystone potential is strongly skewed to the right, with a few taxa showing a relatively high keystone potential when compared to all other taxa (Fig 3A and B, respectively). This aligns well with our understanding of keystones and is to be expected based on the probability density functions of the network features used to compute keystone potential (Fig 2B). Specifically, we observe that the genera *Methanobrevibacter, Agathobaculum, Bilophila* and *Holdemania* (Table 1) have a notably high potential to act as correlation network keystones. We also observe a group of unclassified *Firmicutes* with high keystone potential, but as this group of species is likely not phylogenetically coherent, we did not include it in further analyses. On species level, *Firmicutes bacterium CAG* 83*, Eisenbergiella tayi, Veillonella atypica* and *Ruminococcus lactaris* exhibit a high keystone potential (Table 1). With the exception of the genera *Methanobrevibacter* and *Bilophila*, which belong to the phylum *Euryarchaeota* and *Thermodesulfobacteriota*, respectively, all of these taxa are *Firmicutes*. When we compare the keystone potential of each species with its respective keystone potential on genus level, there is no clear pattern (Fig 3C). This suggests that certain genera may act as a keystone towards other genera but lose this potential on species level, and vice versa. We observe this for *Agathobaculum, Bilophila, Holdemania* and *Methanobrevibacter*. These four genera show a high keystone potential, but the respective species exhibit low keystone potential (Fig 2C). This is a particularly surprising observation considering that only one species belonging to each of these genera is present in the datasets, namely *Agathobaculum butyriciproducens, Bilophila wadsworthia, Holdemania filiformis* and *Methanobrevibacter smithii*. These keystone genera are effectively keystone species that seem to have stronger network keystone properties when considering intergenic correlations. The correlations might be diluted on a species level and intergenic correlations may be distributed amongst multiple species with differing abundance patterns. Conversely, particular species have a high keystone potential that is diluted when observed on genus level, as can be seen for *V. atypica* and *E. tayi.* The species within one genus may rarely co-occur, resulting in opposing correlation patterns and leading to a low keystone potential on genus level. Ultimately, only few taxa in species and genus correlation networks exhibit a high potential to act as keystones, while the vast majority of taxa do not show the characteristics of putative keystones.

**FIG 3.**
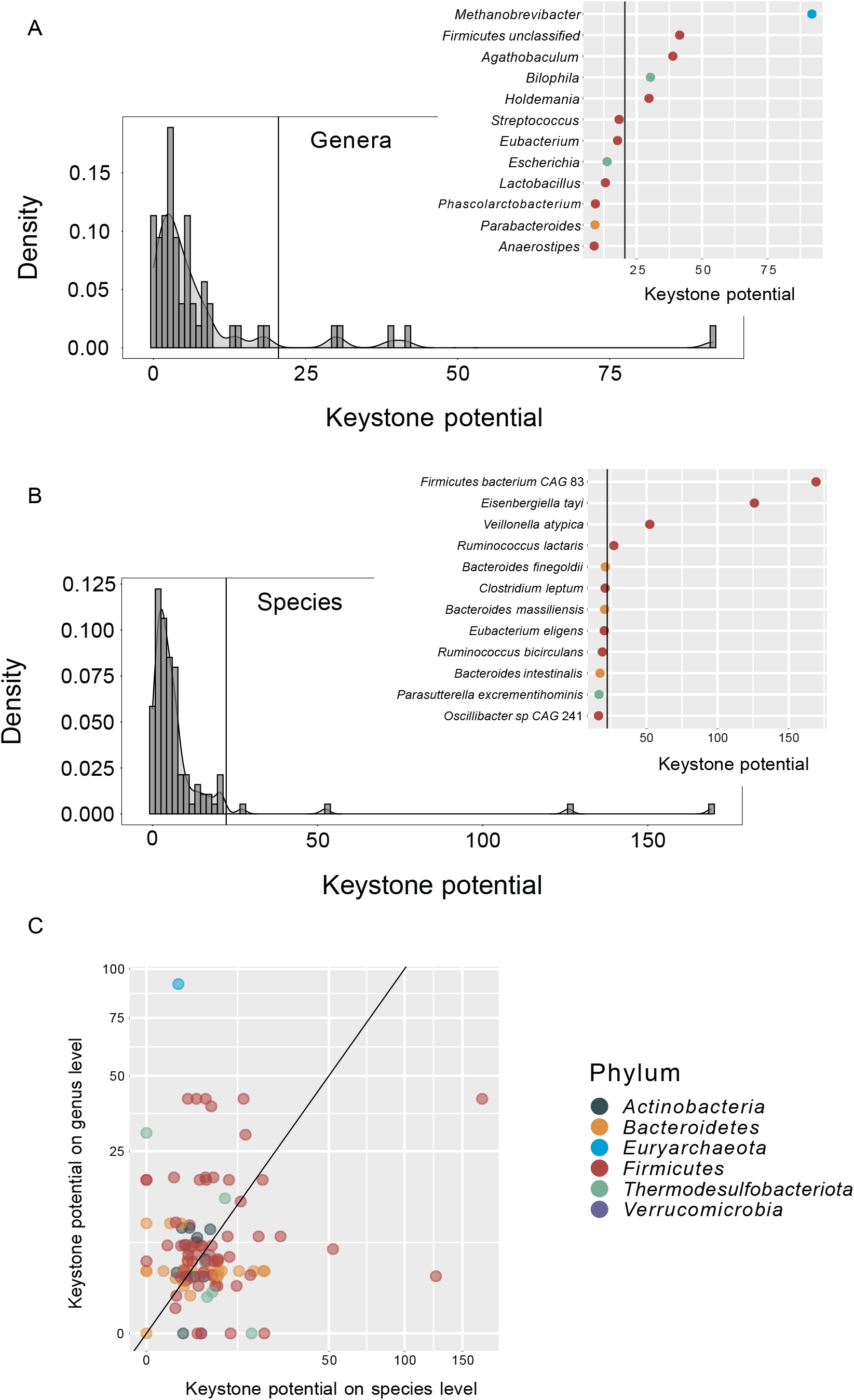
Density functions of keystone potential on (**A**) genus and (**B**) species level. Inserts show the taxa with the highest keystone potential. Horizontal lines indicate the keystone potential cut-off for taxa considered a keystone. The cut-off is set at median + 5x median absolute deviation. (**C**) Keystone potential on species level versus keystone potential on genus level, shown on a logarithmic scale. The diagonal line indicates a 1:1 linear relationship. Colors indicate the phylum of each taxon.

**Table 1.**
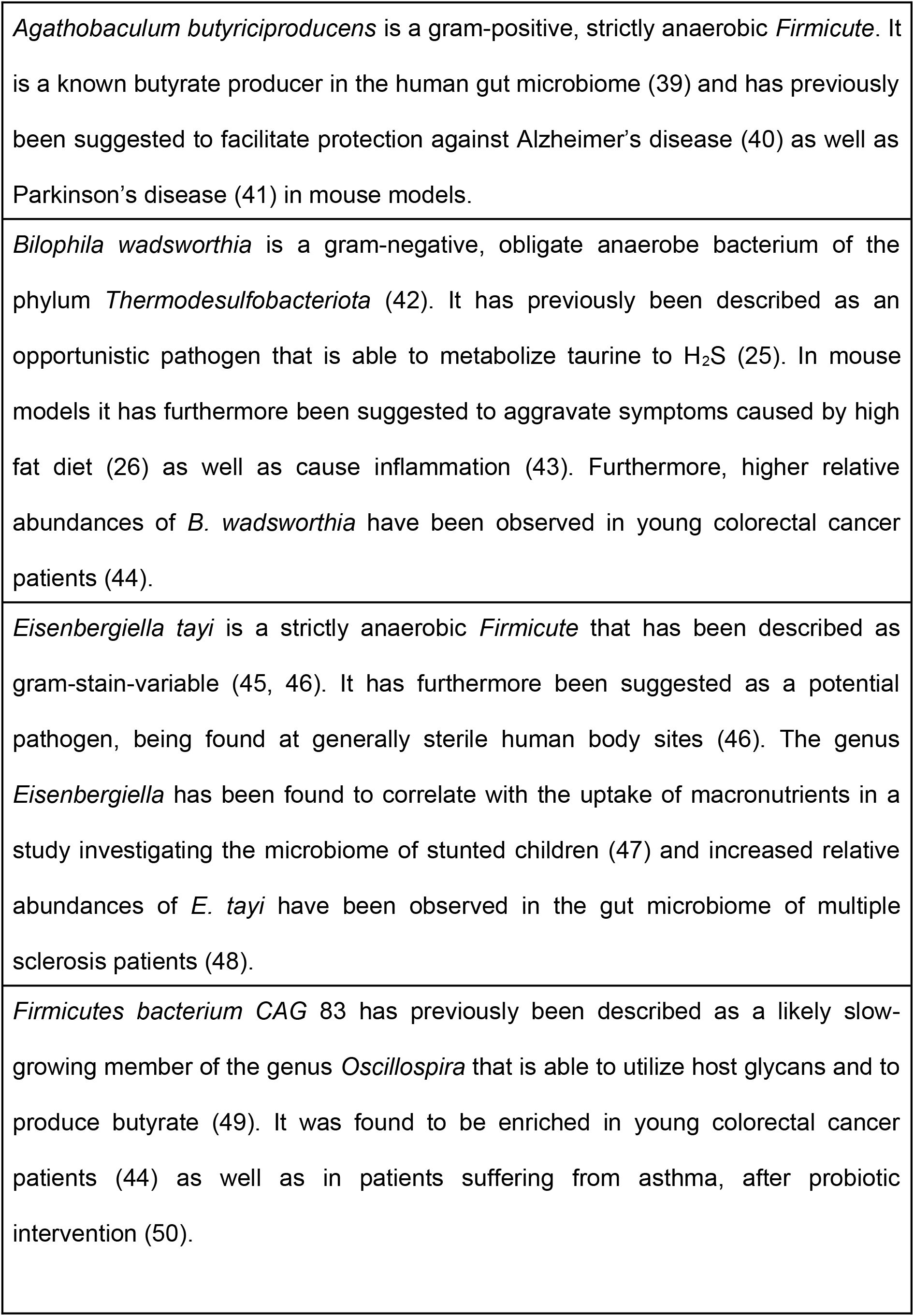
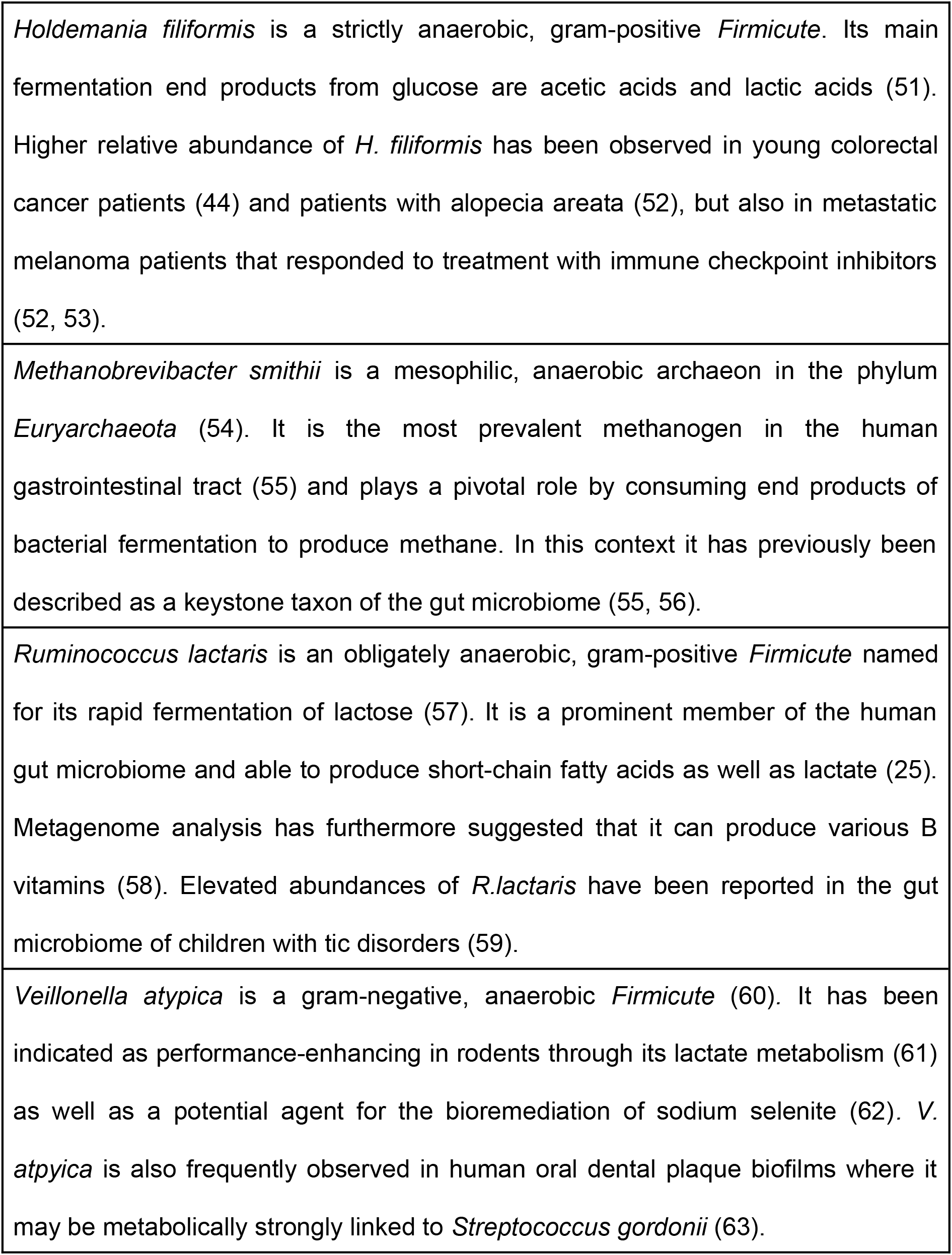
Characteristics of identified correlation network keystone taxa.

### Correlation patterns indicate that keystone taxa may facilitate community stability

It has been suggested that in a highly diverse and functionally redundant system, such as the human gut microbiome, stability is facilitated by a high number of negative interactions, preventing the formation of positive feedback loops (23). However, we observe that more than 50% of correlations in the analyzed networks are positive (Fig 4A). This is even more pronounced in species that belong to the same genus, where almost 75% of observed within-genus correlations are positive (Fig 4B). Putative keystone genera show a contrasting pattern, with around 60% negative correlations (Fig 4C) and genera that are strongly correlated with a keystone exhibit weaker within-genus correlations (Fig 4D). These results suggest that single-species keystone genera might provide community stability by counteracting positive feedback loops within other genera and dampening correlations. We also observe that larger genera exhibit weaker positive correlations within a respective genus (Fig S2A) which may be a result of functional redundancy in these more closely related taxa. On both genus as well as species level we observe that taxa correlated with a keystone taxon (first neighbors of keystones) are in general correlated to more taxa than those not correlated to a keystone. First neighbors of keystones also tend to have weaker correlations (Fig S2B and C, respectively). These observations further point towards a potential stabilizing effect of putative keystone taxa.

**FIG 4.**
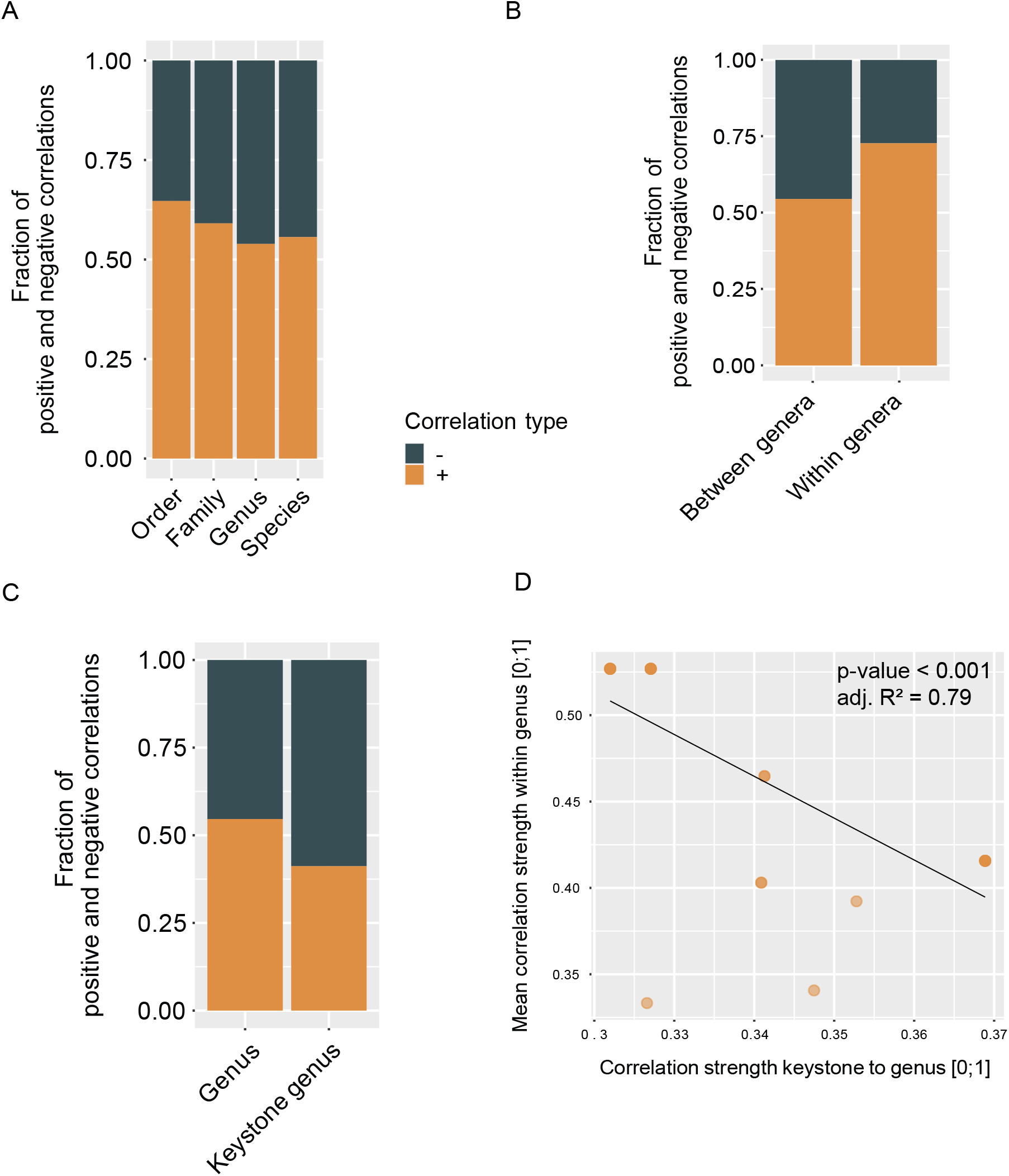
Fractions of negative and positive correlations (**A**) between taxa at different taxonomic resolution, (**B**) between genera and within genera and (**C**) of keystone genera and non-keystone genera. (**D**) Absolute correlation strength from keystone genera to other genera versus the mean absolute correlation strength within the respective genera (Linear model: p-value < 0.001, adj R² = 0.79).

### Correlation patterns reveal co-occurring clusters of taxa

To gain a better understanding of the community assembly characteristics of putative keystone taxa, we investigated the co-occurrence patterns of all prevalent species and genera. We constructed positive correlation networks, taking into account only correlations with a correlation strength > 0, and performed a clustering analysis on these networks. This analysis revealed four distinct clusters of species that are highly positively correlated with each other and that show fewer positive correlations to species of other clusters (Fig 5A). Each keystone species is a member of a different species cluster, indicating that keystone species are not highly correlated with each other. In contrast, keystone genera do not fall into separate clusters in the genus network, despite the presence of again four clusters (Fig 5B). Specifically, *Bilophila* and *Holdemania* are part of one cluster while *Agathobaculum* and *Methanobrevibacter* are located in another. The remaining two clusters are a cluster including *Bacteroides*, a highly diverse and abundant genus, and a cluster including *Prevotella*, a genus with, as previously mentioned, distinct abundance patterns. It should further be noted that high abundance is not a necessity for a correlation network keystone (Fig S1C). In fact, amongst all prevalent taxa (at least 20% prevalence), keystones exhibit medium to low mean relative abundance. Examining the distribution of all network clusters across samples, we observe a gradient of community compositions both on species and genus level (Fig S3). At both taxonomic resolutions there are two dominant clusters that generally comprise the majority of the community. However, all four respective clusters co-occur in most samples and the composition ranges from heavily dominated by one cluster to almost equal contribution of each cluster. This observation is in accordance with recently described entersignatures of gut microbial communities (24) and may point towards a modular gut microbiome composed of distinct groups of co-occurring taxa.

**FIG 5.**
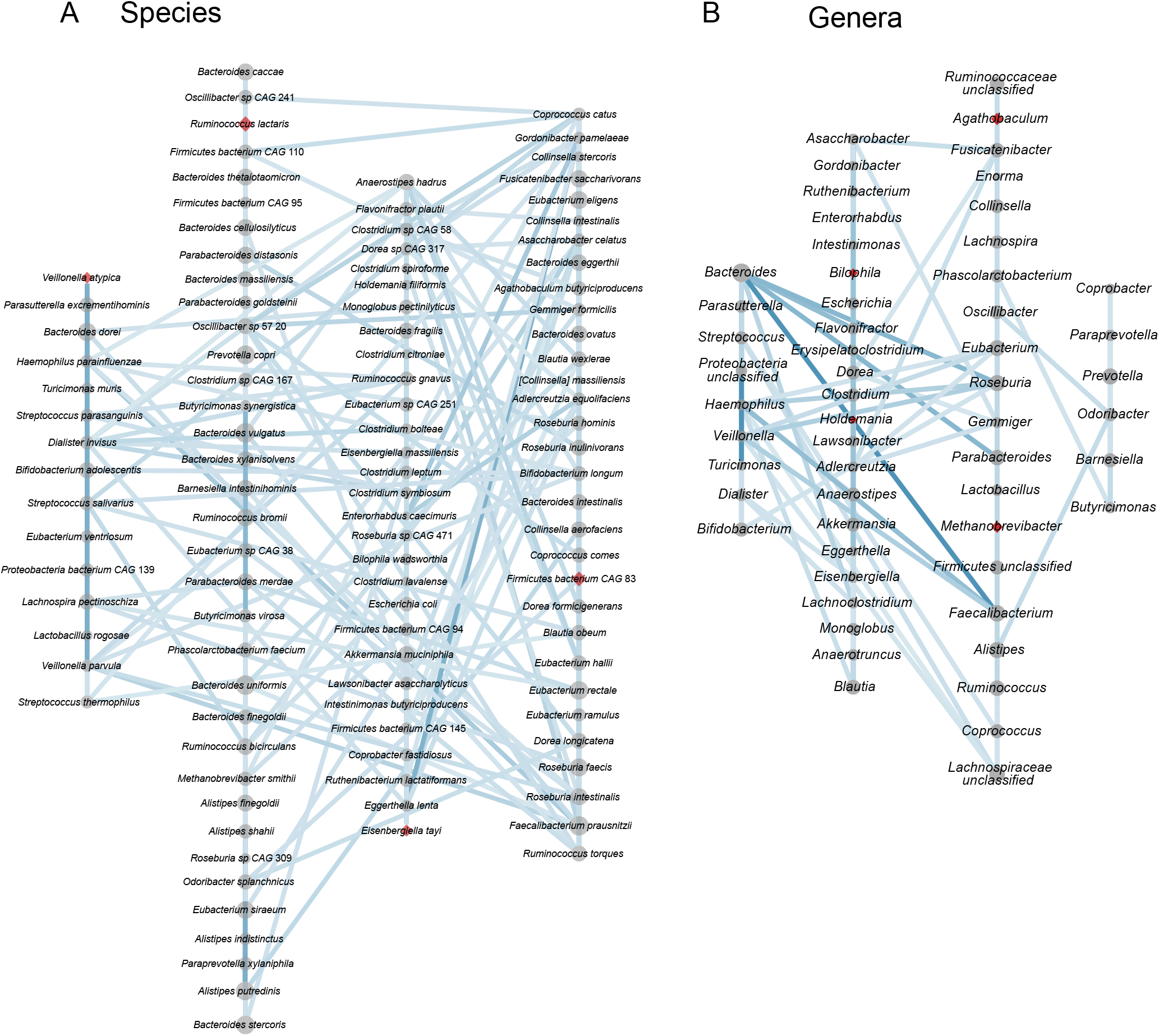
Positive correlation networks of prevalent (present in >= 20% of analyzed samples) (**A**) species and (**B**) genera. Gray circles indicate taxa, red diamonds indicate identified correlation network keystone taxa. Circle and diamond size represents the mean relative abundance of taxa. Edges between circles indicate positive correlations. Light blue indicates a weak correlation, darker blue indicates a stronger correlation. Taxa are organized into 4 clusters of strongly positively correlated species and genera, respectively, as identified with a fast greedy clustering algorithm.

### Keystone taxa are transcriptionally versatile and show distinct transcriptional states

In an effort to elucidate the functional role of putative keystone taxa, we investigated their genomic potential and transcriptional versatility. No genomic features were identified to be distinctive of keystone taxa (Fig S4), so we focused on analyzing their transcriptional versatility and potential transcriptional states. We estimated the transcriptional versatility of prevalent taxa by computing pairwise Sorensen similarities on the presence and absence of gene families in the metatranscriptomes. The average transcriptional similarity of keystone taxa ranges from fairly high (*Holdemania*: 0.7, *R. lactaris:* 0.64) to the lower end of the observed range (*Bilophila*: 0.22, *E.tayi:* 0.18) (Fig S5). However, as is the case for most of the prevalent taxa, transcriptional similarity within each keystone taxon varies strongly across all analyzed samples. This suggests that keystone taxa do not occupy a conserved ecological niche - as inferred by transcriptional profiles - but are rather functionally versatile and potentially occupy different ecological niches within different gut microbial communities. Based on these observations we performed clustering analyses on the transcriptional profiles of the keystone taxa to identify potentially differing transcriptional states. We were able to identify two to three discrete transcriptional states for most keystone taxa (Figs 6, 7), with the exceptions of *Holdemania* and *V. atypica*. In these two cases very few reads mapped to known gene families and fewer still could be grouped into known enzyme commission numbers (ECs), indicating low annotation rates in their reference genomes. Consequently, a clustering analysis based on transcriptional profiles could not be performed for these two taxa.

**FIG 6.**
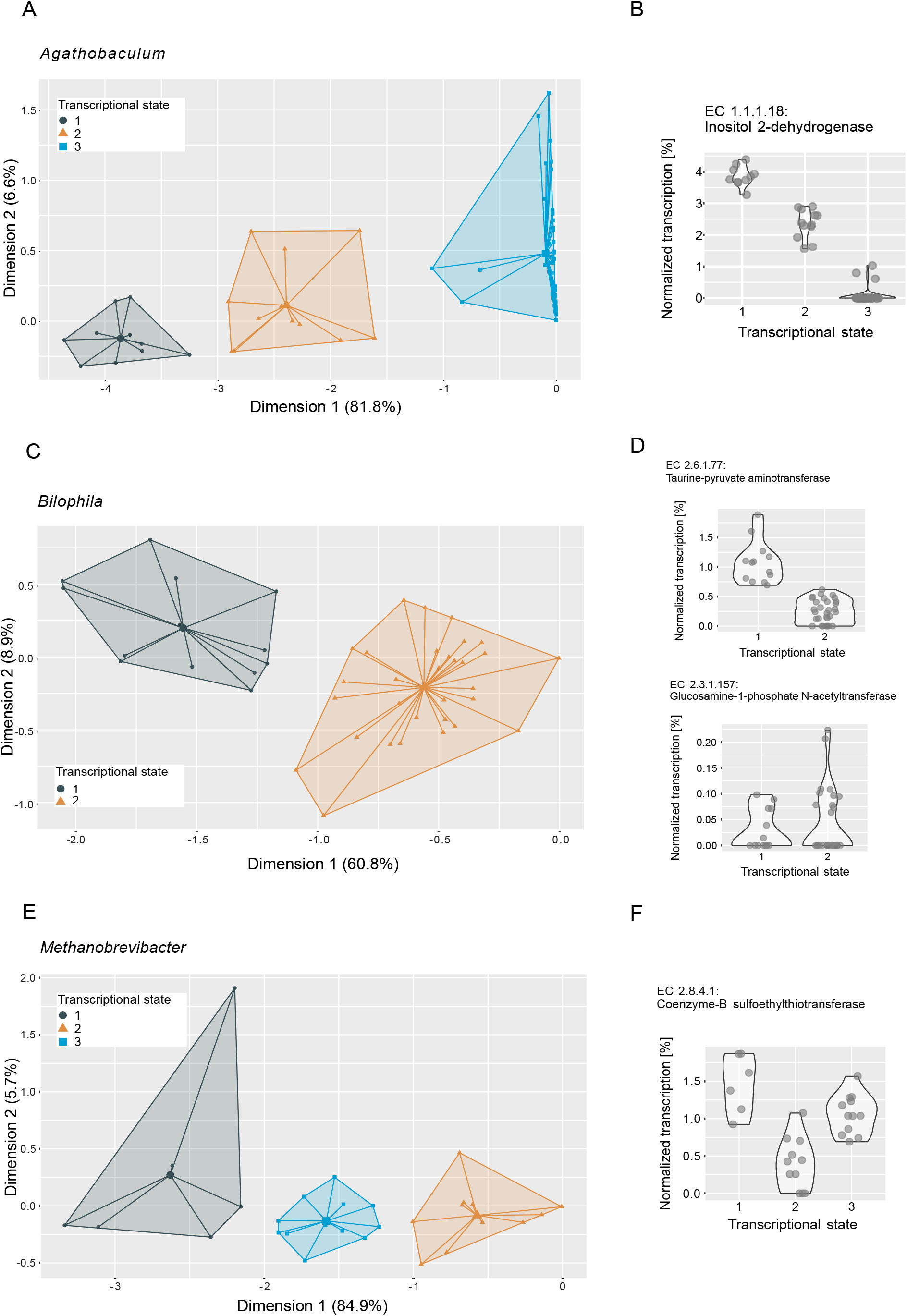
Ordination plots depicting principal component analyses of metatranscriptomes of (**A**) *Agathobaculum*, (**B**) *Bilophila* and (**C**) *Methanobrevibacter.* Metatranscriptome reads were mapped to reference genomes, grouped to Enzyme commission numbers (ECs) and normalized to the total number of reads mapped to each taxon per sample. Colors and icons indicate transcriptional states identified through k-means clustering (average silhouette width: *Agathobaculum:* 0.87, *Bilophila*: 0.92, *Methanobrevibacter*: 0.86). Normalized transcription of example ECs identified as important in distinguishing the transcriptional states through a random forest model are depicted in panels **B, D** and **F**, respectively.

We next performed Random Forest machine-learning analyses to identify individual ECs that distinguish the transcriptional states (Figs 6, 7, Supplementary Table 1). For *Bilophila*, this revealed two ECs with contrasting transcription profiles (Fig 6C and D). Taurine-pyruvate aminotransferase (EC 2.6.1.77), is involved in taurine and hypotaurine metabolism and shows higher relative transcription in the samples in transcriptional state 1. Taurine metabolism is well described in *Bilophila* and may be linked to disease phenotypes (25, 26). Glucosamine-1-phosphate N-acetyltransferase (EC 2.3.1.157), which shows higher relative transcription in transcriptional state 2, is involved in the metabolism of various amino sugars and nucleotide sugars, such as glucose or extracellular N-Acetyl-D-glucosamine. For *Methanobrevibacter*, we could identify three transcriptional states that are likely related to methane metabolism (Fig 6E). Coenzyme-B sulfoethylthiotransferase (EC 2.8.4.1), the enzyme that catalyzes the final step of methanogenesis, was the most important in distinguishing these states (Fig 6F). Our analysis revealed three additional ECs involved in methane metabolism (EC 2.3.1.101, Formylmethanofuran-tetrahydromethanopterin formyltransferase, EC 1.17.1.9, Formate dehydrogenase and EC 4.1.2.43, 3-hexulose-6-phosphate synthase) as important in distinguishing transcriptional states (Table S1). Two transcriptional states were identified in *E. tayi* (Fig 7C). Notably, all three ECs identified as important in distinguishing these transcriptional states are involved in the biosynthesis of lysine (Fig 7D). The proportion of *E. tayi* transcription dedicated to these enzymes seems to be higher in transcriptional state 2, pointing towards an upregulation of lysine biosynthesis. It is important to note that a large proportion of metatranscriptome reads could not be grouped into any known ECs. We chose the higher resolution of ECs over gene families to gain more insight into the function of the transcripts at the expense of losing a larger proportion of reads. Despite these limitations, the data point towards the presence of functionally versatile keystone taxa in the gut microbial community.

**FIG 7.**
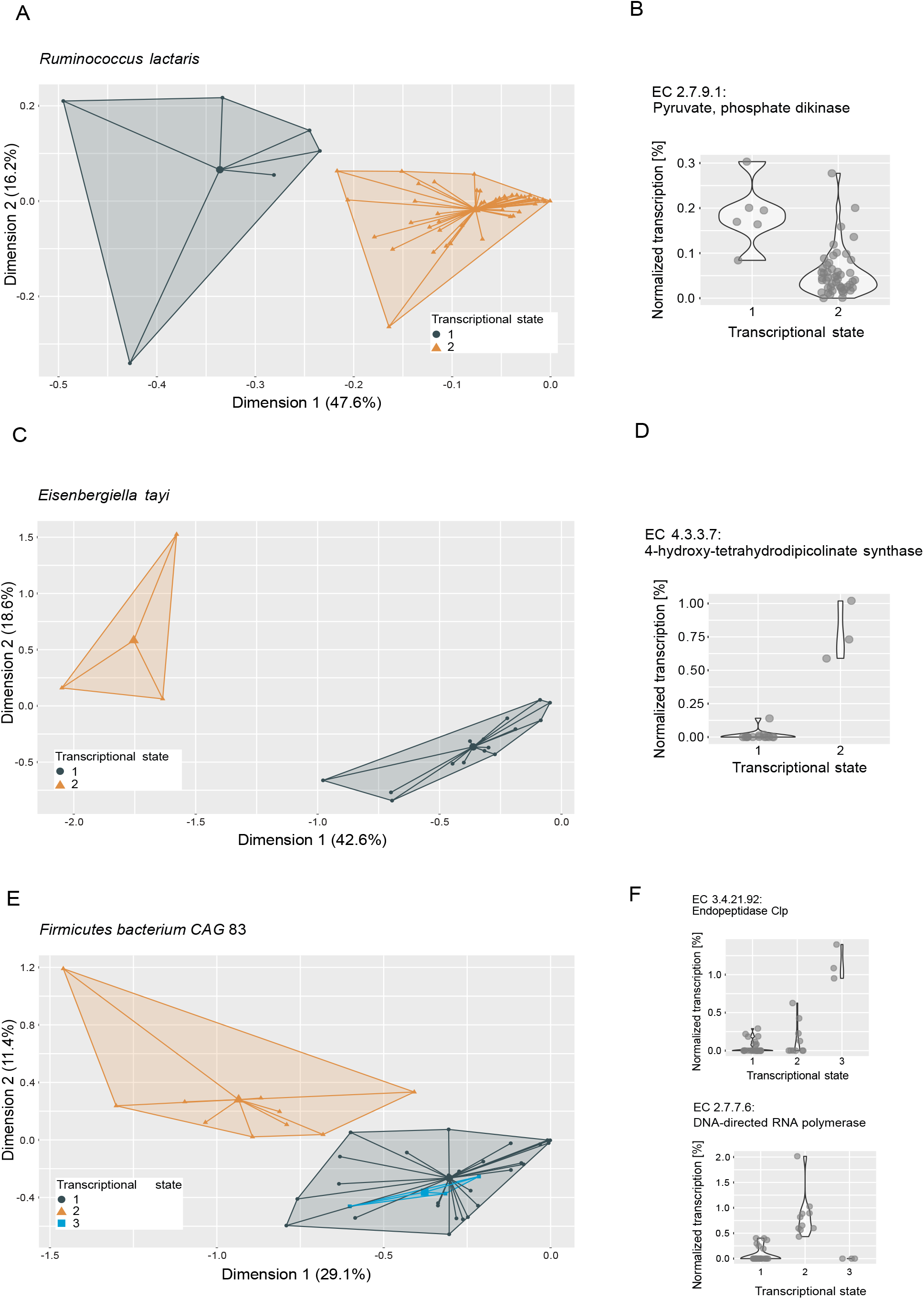
Ordination plots depicting principal component analyses of metatranscriptomes of (**A**) *Ruminococcus lactaris*, (**B**) *Eisenbergiella tayi* and (**C**) *Firmicutes bacterium CAG* 83. Metatranscriptome reads were mapped to reference genomes, grouped to Enzyme commission numbers (ECs) and normalized to the total number of reads mapped to each taxon per sample. Colors and icons indicate transcriptional states identified through k-means clustering (average silhouette width: *R.lactaris:* 0.92, *E.tayi*: 0.78, *Firmicutes bacterium CAG* 83: 0.81). Normalized transcription of example ECs identified as important in distinguishing the transcriptional states through a random forest model are depicted in panels **B, D** and **F**, respectively.

## DISCUSSION

The aim of the present study is to elucidate the potential ecological role of correlation network keystone taxa of the human gut microbiome and gain a better understanding of how these analyses tie into other avenues of human gut microbiome research. In order to identify correlation network keystones, we developed a workflow based on a bootstrapping approach and a subsampling regime that enables the construction of robust and statistically significant correlation networks. We performed network analysis at the taxonomic resolutions of order, family, genus and species and calculated the keystone potential of prevalent taxa combining their mean degree, transitivity and betweenness centrality. We only observe high keystone potential of any taxa in genus- and species-level networks (Fig 2C), in contrast to previous suggestions of higher-order keystones (11). While we were able to identify taxa with a high keystone potential in the genus-level correlation network, they are interestingly all single-species genera, meaning that no other species belonging to these genera were detected in the analyzed samples. We therefore only identify putative keystone species rather than genera. A lower taxonomic resolution inevitably increases the functional potential of a taxon and broadens its ecological niche, which thereby likely weakens direct and indirect interactions between taxa. This would explain the more evenly distributed correlations between orders and families when compared to genera and species and the lower keystone potentials. Keystone taxa are of great interest for systematic microbiome engineering, for example for the restoration of a dysbiotic gut microbiome through the introduction or targeted manipulation of keystones. In applied fields a microbial species or genus may well present a more practical target, as opposed to broader taxonomic groups.

Overall, the correlation networks have a high percentage of positive correlations (Fig 4A). This is somewhat surprising, as work by Coyte and colleagues (23) has suggested that a diverse microbial community can only be stable when the majority of interactions are negative. Interestingly, the single-species keystone genera identified through our analysis have a considerably higher percentage of negative correlations to other genera (Fig 4C). These high numbers of negative correlations with keystone taxa may provide stability to the microbial community, especially considering the particularly high percentage of positive correlations between species of the same genus (Fig 4B). The negative correlations with keystone genera potentially counteract these positive correlations that could otherwise result in positive feedback loops and lead to instability. Coyte and colleagues (23) identify the dampening of these positive feedback loops as one mechanism for stabilization of a community. They furthermore suggest weaker ecological interactions as an additional stabilization mechanism. We indeed observe that the stronger the correlation of a keystone to a genus, the weaker the correlations between the species of the respective genus (Fig 4D), suggesting that the keystone genera may be both dampening positive feedback loops and weakening intergenic correlations. We additionally find that species’ positive correlations tend to be weaker in larger genera (Fig S2A). This result supports another observation by Coyte et al. (23) that functional redundancy in closely related taxa provides stability by replacing few strong interactions with many weaker interactions. In general, taxa correlated to a keystone taxon tend to have more, but weaker correlations (Fig S2B and C), again indicating a stabilizing effect from keystone taxa.

In both the species- as well as genus-level network we identified four clusters of co-occurring taxa (Fig 5). Each of the four species-clusters contains exactly one putative keystone species, while only two of the genus-clusters contain two keystone genera each. These respective four clusters are in turn frequently co-occurring (Fig S3). The analyzed gut microbiomes all consisted of more than one cluster, both on species as well as genus level, resulting in a compositional gradient ranging from strongly dominated by a particular cluster to a fairly even distribution of all four clusters. Frioux and colleagues (24) recently described similar patterns in the genus composition of gut microbial communities. They identified five so-called enterosignatures (commonly co-occurring genera) that, in combination, can accurately describe most healthy gut microbiomes. These five enterosignatures show strong similarities to the genus clusters we were able to identify in our correlation network. Specifically, ES-Firm (enterosignature with a high contribution of various *Firmicutes*) and ES-Prev (dominated by *Prevotella*) correspond to cluster 3 and cluster 4 in Figure 4, respectively. An enterosignature mainly found in the gut microbiome of infants, ES-Bifi, does not have a corresponding genus cluster in the cohort we analyzed, which consists solely of adults. However, the main genera of ES-Bifi, namely *Bifidobacterium, Streptococcus, Veillonella, Enterococcus* and *Haemophilus*, are members of genus cluster 1, together with *Bacteroides*, one of the main genera in ES-Bact. The remaining cluster 2 includes *Escherichia*, the main contributor to ES-Esch, and *Blautia*, a strong contributor to ES-Firm, but most genera in this cluster were not reported as strongly contributing to an enterosignature. These differences may be due to differences in the applied methods (correlation analysis vs. non-negative matrix factorization) as well as size and diversity of the analyzed cohorts. While not a perfect match to the described enterosignatures, the genus clusters we identified are still remarkably similar in composition.

We observe distinct transcriptional states in all putative keystone taxa that can be attributed to differential transcription of particular ECs. In a small synthetic gut microbial community, Shetty and colleagues (27) observed shifting functional roles of species in an otherwise compositionally and functionally fairly stable community. These observations suggest functional versatility that enables these taxa to adjust their metabolism to available resources and shifting ecological niches. Particularly striking is the high number of ECs from *Methanobrevibacter* involved in methane metabolism that we could identify as important in distinguishing the transcriptional states of *Methanobrevibacter.* In *Bilophila,* we identified an EC involved in taurine and hypotaurine metabolism as important in distinguishing transcriptional states and an EC potentially involved in host glycan degradation that shows an opposing transcription pattern, suggesting a metabolic shift. These findings are consistent with the idea that the gut microbiome needs to be able to accommodate a diverse and variable set of available resources as well as dynamic interplay with the host.

Together with the stabilizing effects previously discussed, functional versatility may be another mechanism enabling keystone taxa to exert their influence on the microbial community.

## LIMITATIONS OF THE STUDY

The present study has several limitations. We focused exclusively on the analysis of publicly available data and no experimental validation was carried out. Metagenome and metatranscriptome reads were mapped to reference genomes and analyzed on different taxonomic resolutions up to species. Consequently, strain-level variation has not been taken into account and the reliance on reference genomes further reduces the genetic variation represented in this study. We also acknowledge that correlation networks can reflect other ecological processes beyond interactions such as habitat filtering. Additionally, our analyses focused only on prevalent taxa and therefore would miss any less common keystone taxa. Our observations are furthermore limited to the distal colon, while community composition as well as functionality may differ in more proximal parts of the intestines. Lastly, the data we used for this study consisted of two cohorts with limited demographic diversity and different putative keystones may be present in other populations.

## CONCLUSION

In this study we developed a pipeline for the robust construction and analysis of correlation networks and the identification of putative keystone taxa. Through a comprehensive analysis of the constructed correlation networks we are able to show that correlation networks and their properties are highly sensitive to taxonomic resolution. Furthermore, we identified signatures of potentially community stabilizing interaction patterns reflected in the correlation networks as well as co-occurring sub-communities in the human gut microbiome that show a high similarity to previously described enterosignatures. The ecological significance of correlation network keystones is still an open question. Nevertheless, we see indications that keystone taxa may have a local stabilizing effect on sub-communities of the gut microbiome. This suggests that it is unlikely that there are individual taxa that globally affect the entire community. We rather hypothesize that keystone taxa act as stabilizing agents in relatively independently co-occurring subsets of the microbiome.

## MATERIALS AND METHODS

### Processing of raw data

We analyzed metagenome and metatranscriptome reads from two publicly available datasets, a cohort of patients with inflammatory bowel disease as well as healthy control patients (18) and a cohort of adult men (19). The raw reads were preprocessed with trimmomatic (28) (with settings leading: 3, trailing: 3, slidingwindow: 4:15, minlen: 50). We removed samples with fewer than one million reads from further analyses and, in the Schirmer dataset, used only samples from participants not diagnosed with IBD. We then randomly selected one sample per participant to ensure statistical independence of the samples. The trimmed reads were processed using HUMAnN 3 (29). Briefly, HUMAnN 3 first estimates community composition with MetaphlAn 3 (29), second it maps reads to a community pangenome with bowtie2 (30) and third it aligns unmapped reads to a protein database using DIAMOND (31). This results in both taxonomic as well as functional profiles of the metagenome and metatranscriptome reads. We used the setting rel_ab_w_read_stats in MetaphlAn 3 to estimate both relative abundances and total reads mapped to a reference genome. In the functional profiles we further grouped reads mapped to gene families into enzyme commission numbers (ECs) using the HUMAnN 3 function humann_regroup_table with - -groups uniref90_level4ec.

### Principal coordinate analysis of community profiles

In order to visualize community similarity and overlap between the two analyzed datasets, we computed the robust Aitchison distance between samples with the function vegandist in the R package vegan (32), using the total estimated reads output from MetaphlAn 3 on species resolution. We then performed a principal coordinate analysis on these distances using the function cmdscale in the R package stats (33).

### Computation of correlation networks

We computed binary Jaccard similarities of community composition at species resolution between samples to estimate whether the communities were similar enough to compute correlation networks. Berry and Widder (13) suggest an overall site similarity (Jaccard similarity) of at least 20% and we observed a mean Jaccard similarity of 36%. We computed correlations between taxa using FastSpar (16) with the settings - -iterations 100 (corresponds to rounds of SparCC correlation estimations), - -exclude_iterations 20 (number of times highly correlated taxa pairs are discovered and excluded), - -threshold 0.1 (minimum threshold to exclude correlated taxa pairs) and - -number 1000 (number of bootstraps). FastSpar is a C++ implementation of the correlation network analysis algorithm SparCC (17). In short, these algorithms use log-transformed components to estimate linear Pearson correlations to account for the compositional nature of metagenomic data. In order to estimate the significance of observed correlations, FastSpar uses a permutation based approach to generate a null distribution that observed correlations are then compared against. We used the total estimated reads output from MetaphlAn 3 (29) as abundance estimates for each taxon. We computed correlations between taxa at the taxonomic resolution of order, family, genus and species. To ensure robust results, we subsampled our combined datasets. Specifically, we used FastSpar to compute correlation networks on a random subsample of 50 samples and repeated this process 1000 times. We then built a consensus network using only correlations with a p-value <= 0.05 in at least 200 iterations. The correlation strength in the consensus network was estimated as the mean correlation strength of significant correlations across all iterations. This workflow is visualized in Figure 1. We calculated network features, namely modularity, cohesion, relative node degree, closeness centrality, betweenness centrality and transitivity, using the R package igraph (34). Following the suggestion of Berry and Widder (13) we estimated each taxon’s potential to act as a keystone using the network features relative node degree, betweenness centrality and transitivity. We excluded closeness centrality from our calculations in order to reduce collinearity of features as we saw strong linear relationships between relative node degree and closeness centrality at all taxonomic resolutions (data not shown). As shown by Berry and Widder (13), correlation network keystone taxa show a high relative node degree and transitivity as well as low betweenness centrality. We therefore computed the keystone potential of each taxon as (relative node degree * transitivity) / betweenness centrality. We consider taxa a keystone if they show a particularly high keystone potential as compared to all other prevalent taxa (taxa present in at least 20% of samples). Namely, we chose a cut-off of median + 5x median absolute deviation of keystone potential after visually examining the entire distribution. We furthermore grouped the prevalent taxa into clusters using the function cluster_fast_greedy with default settings in the R package igraph (34) on positive correlation networks, in which only correlations with a correlation strength > 0 are considered. We visualized correlation networks using Cytoscape version 3.8.0 (35).

### Functional potential of keystone taxa

We investigated the functional potential of keystone taxa using the pathabundances output from HUMAnN 3 (29). In short, pathway abundances are computed on community level as well as for individual taxa based on the abundances of the individual component reactions constituting an entire pathway. The pathway definitions used are based on MetaCyc (36).

### Transcriptional stability and transcriptional states of keystone taxa

In order to estimate functional stability of taxa we computed pairwise Sorenson similarities on the presence and absence of gene families using the function betadiver in the R package vegan (32). We performed k-means clustering to further analyze the functional profiles of the keystone taxa and identify different transcriptional states. For this analysis we opted to use the functional resolution of ECs provided by HUMAnN 3 (29) (function humann_regroup_table with - - groups uniref90_level4ec) and normalized copies per million to the total transcription attributed to each taxon per sample. We use the function fviz_nbclust in the R package factoextra (37) to choose the optimal number of clusters (with a k.max - maximum number of clusters - of 10) by computing the average silhouette width for each number of clusters and choosing the highest score. For this number of clusters we then performed k-means clustering using the function kmeans in the R packages stats (33). However, if we observed clusters containing only 1 or 2 samples, we excluded these samples from the analysis and repeated the process. In order to gain some insights into which ECs are particularly important for distinguishing the observed transcriptional states, we performed random forest analyses. A random forest analysis is a classifying algorithm based on decision trees that calculates importance metrics for the features used to cluster a dataset. In order to estimate the significance of these importance metrics we used the function rfPermute in the R package rfPermute (38). This function computes a null distribution of importance metrics by permuting the response variable and calculates p-values for the observed importances. We used the mean decrease in accuracy as the importance metric for our analysis.

## CODE AVAILABILITY

R Code for the established workflow to construct correlation networks and compute keystone potential of individual taxa can be found here: https://github.com/fbauchinger/correlation.network.keystones_workflow

## ACKNOWLEDGEMENTS

This work was supported by ERC starting grant 714623 “FunKeyGut”. Work in the laboratory of DB is supported by Austrian Science Fund project MAINTAIN DOC 69 doc.fund.

DB and FB designed the study. FB and DS performed analyses. DB, FB and DS interpreted the results and wrote the manuscript.

The authors declare that no competing interests exist.

---

**Table S1.** Enzyme commission numbers (ECs) identified as important in distinguishing transcriptional states of keystone taxa^a^.

| **Taxon** | **EC** | **Mean decrease accuracy** | **Involved in the following pathways (64)** |
| --- | --- | --- | --- |
| *Agathobaculum* | 1.1.1.18: Inositol 2-dehydrogenase | 6.870 | Streptomycin biosynthesis<br>Inositol phosphate metabolism |
| *Agathobaculum* | 2.7.13.3: Histidine kinase | 3.204 | - |
| *Agathobaculum* | 1.3.8.1: Short-chain acyl-CoA dehydrogenase | 2.036 | - |
| *Agathobaculum* | 5.3.1.9: Glucose-6-phosphate isomerase | 2.028 | Glycolysis/Gluconeogenesis<br>Pentose phosphate pathway<br>Starch and sucrose metabolism<br>Amino sugar and nucleotide sugar metabolism |
| *Agathobaculum* | 2.4.2.3: Uridine phosphorylase | 1.005 | Pyrimidine metabolism |
| *Bilophila* | 2.6.1.77: Taurine-pyruvate aminotransferase | 4.049 | Taurine and hypotaurine metabolism |
| *Bilophila* | 2.3.1.157: Glucosamine-1-phosphate N-acetyltransferase | 1.754 | Amino sugar and nucleotide sugar metabolism |
| *Bilophila* | 7.1.3.1: H<sup>+</sup>-exporting disphosphatase | 1.604 | - |
| *Bilophila* | 4.1.99.12: 3,4-dihydroxy-2-butanone-4-phosphate synthase | 1.005 | Riboflavin metabolism |
| *Bilophila* | 2.7.4.9: dTMP kinase | 1.005 | Pyrimidine metabolism |
| *Eisenbergiella tayi* | 1.17.1.8: 4-hydroxy-tetrahydrodipicolinate reductase | 1.742 | Monobactam biosynthesis<br>Lysine biosynthesis |
| *Eisenbergiella tayi* | 4.3.3.7: 4-hydroxy-tetrahydrodipicolinate synthase | 1.477 | Monobactam biosynthesis<br>Lysine biosynthesis |
| *Eisenbergiella tayi* | 2.7.2.4: Aspartate kinase | 1.005 | Glycine, serine and threonine metabolism<br>Monobactam biosynthesis<br>Cysteine and methionine metabolism<br>Lysine biosynthesis |
| *Firmicutes*<br>*bacterium* CAG 83 | EC 2.7.7.6: DNA-directed RNA polymerase | 6.068 | - |
| *Firmicutes*<br>*bacterium* CAG 83 | 3.4.21.92: Endopeptidase Clp | 4.772 | - |
| *Methanobrevibacter* | 2.8.4.1: coenzyme-B<br>sulfoethylthiotransfer<br>ase | 2.937 | Methane metabolism |
| *Methanobrevibacter* | 2.3.1.101:<br>formylmethanofuran-<br>tetrahydromethanopt<br>erin N-formyltransferase | 2.291 | Methane metabolism |
| *Methanobrevibacter* | 1.17.1.9: formate<br>dehydrogenase | 1.903 | Glyoxylate and<br>dicarboxylate metabolism<br>Methane metabolism |
| *Methanobrevibacter* | 4.1.2.43: 3-hexulose-6-phosphate<br>synthase | 1.722 | Pentose phosphate pathway<br>Methane metabolism |
| *Methanobrevibacter* | 1.1.1.1: alcohol<br>dehydrogenase | 1.722 | Glycolysis/Gluconeogenesis<br>Fatty acid degradation<br>Glycine, serine and<br>threonine metabolism<br>Tyrosine metabolism<br>alpha-Linolenic acid<br>metabolism<br>Pyruvate metabolism<br>Chloroalkane and<br>chloroalkene degradation<br>Naphthalene degradation<br>Retinol metabolism |
|  |  |  | Metabolism of xenobiotics<br>by cytochrome P450<br>Drug metabolism -<br>cytochrome P450 |
| *Methanobrevibacter* | 2.3.1.174: 3-oxoadipyl-CoA<br>thiolase | 1.719 | Phenylalanine metabolism<br>Benzoate degradation |
| *Methanobrevibacter* | 1.1.1.261: sn-glycerol-1-phosphate<br>dehydrogenase | 1.005 | Glycerophospholipid<br>metabolism |
| *Ruminococcus*<br>*lactaris* | 2.7.9.1: (pyruvate,<br>phosphate dikinase) | 2.276 | Glycolysis/Gluconeogenesis<br>Pyruvate metabolism<br>Carbon fixation pathways in<br>prokaryotes |

<sup>a</sup> Shown are significant predictors (p-value $\leq 0.05$ ). Mean decrease in accuracy is scaled through division by standard error.

**FIG S1.**
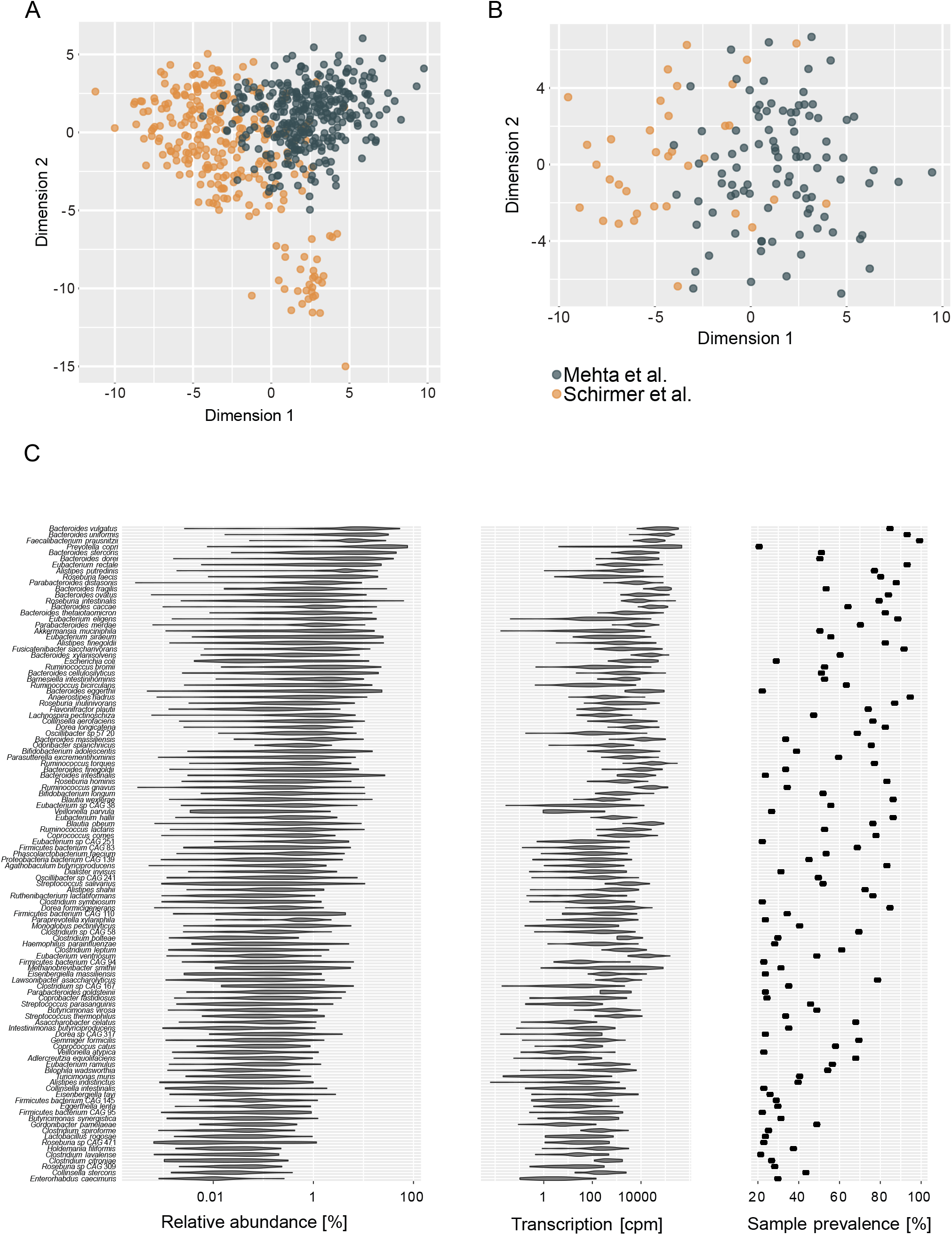
Principal coordinate analysis of (**A**) all processed samples and (**B**) samples used for further analysis (one randomly selected sample per participant). Depicted is the robust Aitchison distance of estimated total reads mapped to species reference genomes. Colors indicate the study that published the raw data. Panel **C** shows relative abundance [%], transcription [cpm - copies per million] and sample prevalence [%] of species present in at least 20% of analyzed samples. Species are ordered by mean relative abundance.

**FIG S2.**
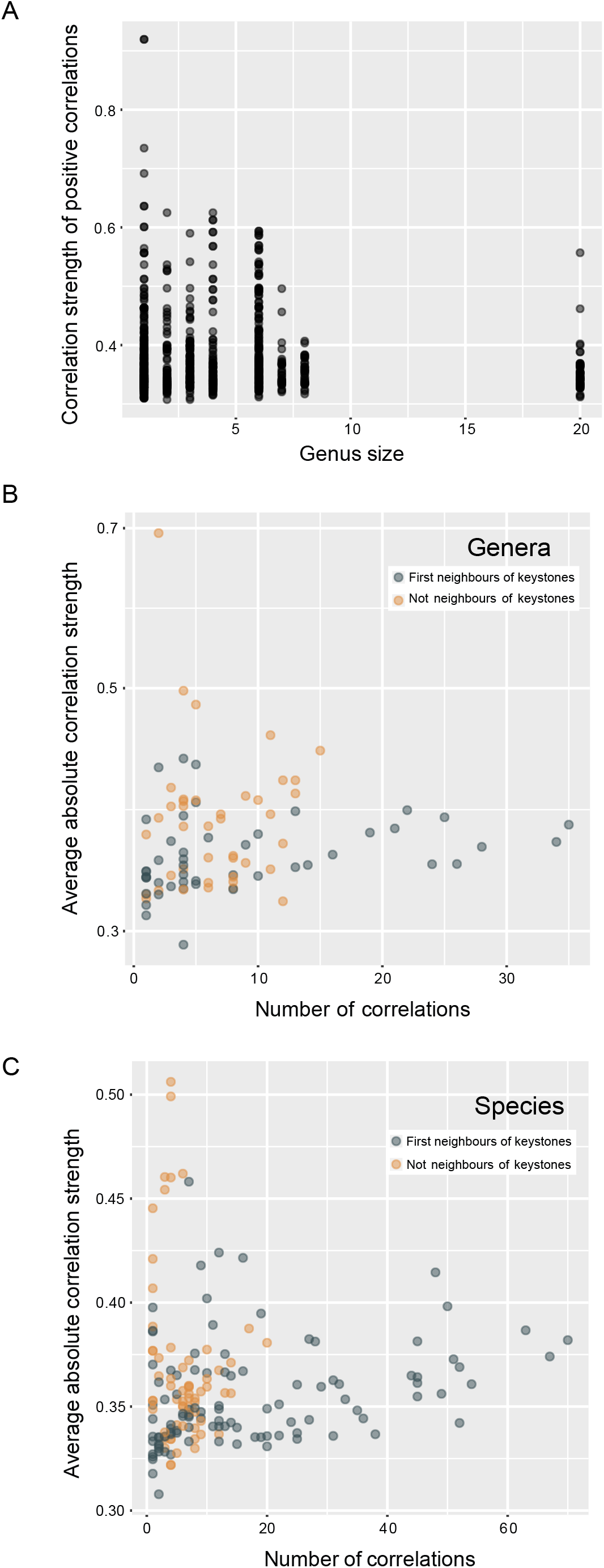
Distributions of network correlations in (**A,C**) the species-level network and (**B**) the genus-level network. Panel **A** shows the within-genus correlation strength of positive correlations as a factor of genus size. Panels **B** and **C** depict the average absolute correlation strength compared to the total number of correlations in (**B**) the genus-level network and (**C**) the species-level network. Gray circles indicate taxa that are first neighbors of (= directly correlated with) a keystone taxon, orange circles indicate taxa that are not first neighbors of any keystone taxa.

**FIG S3.**
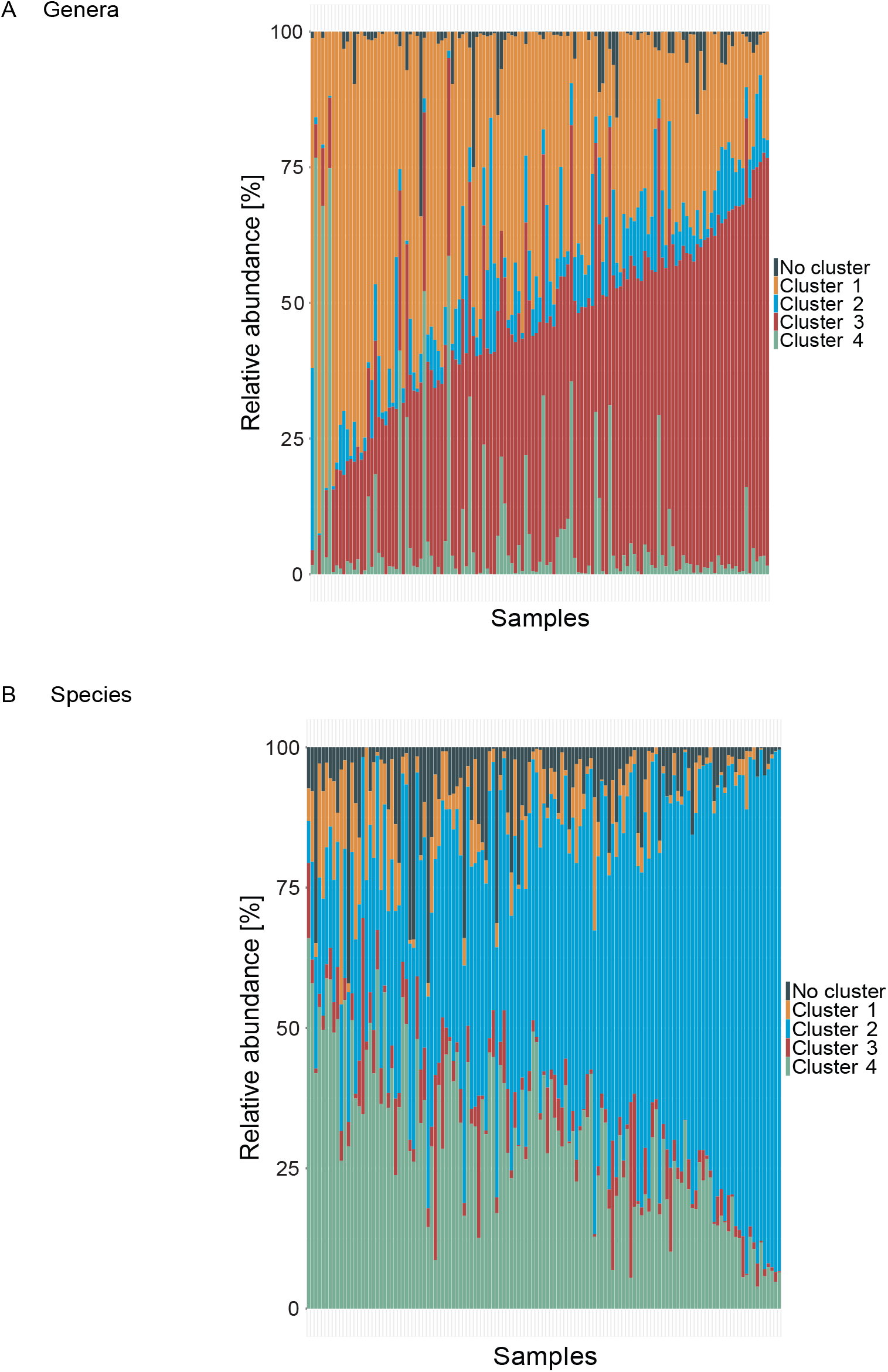
Relative abundance [%] of co-occurrence network clusters of (**A**) genera and (**B**) species. Colors indicate the respective cluster that taxa were grouped into (Fig 5), gray indicates taxa not grouped into any cluster. Samples were ordered based on a k-means clustering algorithm.

**FIG S4.**
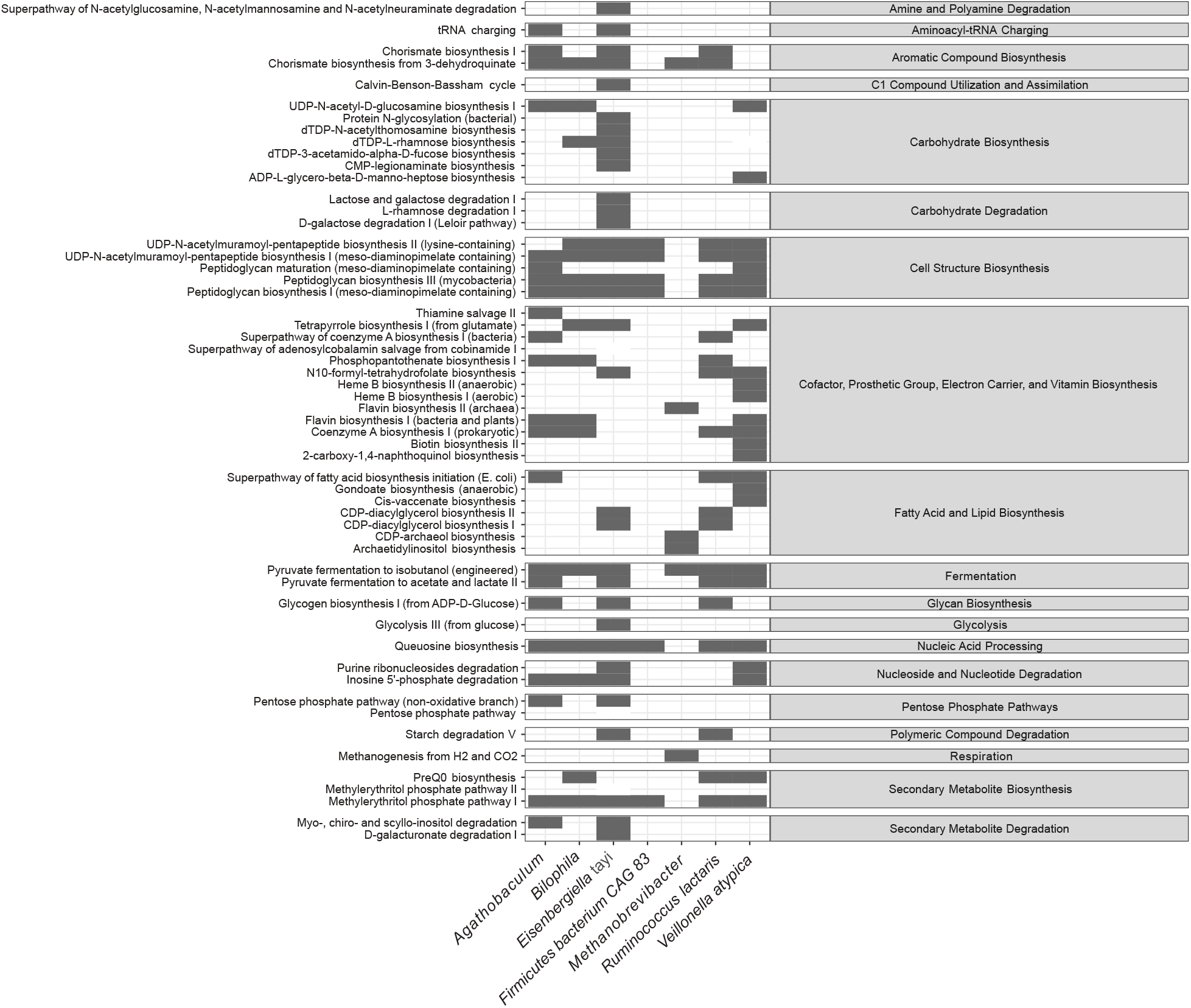
Presence of metabolic pathways in identified correlation network keystone taxa. Gray color indicates that a pathway was found to be present in the metagenome reads mapped to the respective taxon.

**FIG S5.**
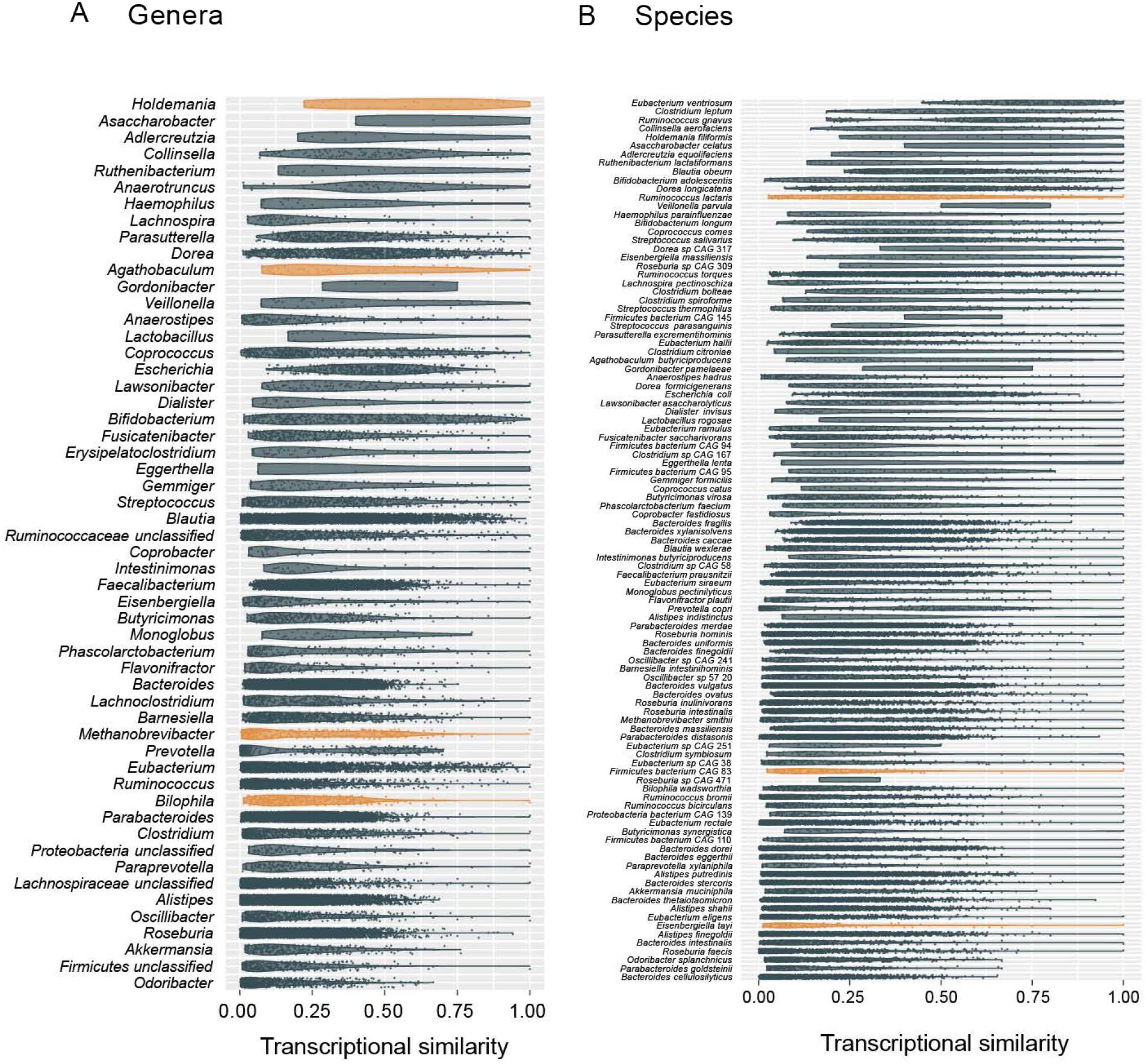
Transcriptional similarity of (**A**) genera and (**B**) species, with 1 indicating identical transcriptional profiles and 0 indicating no overlap. Transcriptional similarity was calculated as pairwise Sorenson distance from the presence and absence of gene families in the metatranscriptomes. Orange color marks the identified keystone taxa. Taxa are ordered by mean Sorenson distance.

